# Predictive Approach to Understanding *Angiostrongylus cantonensis* Distribution in the Canary Islands

**DOI:** 10.1101/2025.11.28.691155

**Authors:** Lucia Anettová, Jan Divíšek, Radovan Coufal, Anna Šipková, Jana Kačmaříková, Michal Horsák, Vojtech Baláž, Elena Izquierdo-Rodriguez, Barbora Červená, Pilar Foronda, David Modrý

## Abstract

*Angiostrongylus cantonensis* is an invasive parasitic nematode and zoonotic pathogen responsible for eosinophilic meningitis. Originally native to Southeast Asia, it is now globally distributed across tropical and subtropical regions and is approaching Europe, with Tenerife as a key hotspot.

This study investigates the distribution and prevalence of *A. cantonensis* in Tenerife across three host groups (rats, gastropods, lizards). Based on prevalence data, we modelled its potential distribution using species distribution models (SDMs) and compared climatic conditions with Hawaii, a region with frequent human cases.

Field surveys confirmed *A. cantonensis* in endemic and introduced gastropods (25.6%; 179/698), rats (21.5%; 14/79), and lizards (24.0%; 31/129), with local prevalence ranging from 2.4% to 41.6%. MaxEnt and Boosted Regression Tree models identified precipitation seasonality as the main driver of distribution, while prevalence was influenced primarily by tree cover density and climatic variability. Northeastern Tenerife, La Gomera, La Palma, and El Hierro showed the highest habitat suitability. However, overlap with densely populated areas was limited, possibly explaining the absence of reported human cases. The MESS analysis, based on climatic data from Hawaii, indicated moderate to high environmental similarity across most of the Canary Islands, except in northeastern Tenerife, where conditions were outside the range observed in Hawaii.

*A. cantonensis* is firmly established in Tenerife, but human cases remain absent, likely due to limited human exposure, cultural practices, and geographic separation of parasite hotspots from urban zones. Our findings highlight the importance of integrating ecological and epidemiological data in zoonotic risk assessments.

**Author Summary:** The rat lungworm, *Angiostrongylus cantonensis*, is a parasitic nematode that can infect the human brain and cause a serious disease known as eosinophilic meningitis. Although it was once limited to Southeast Asia, it has now spread across many tropical regions, and its arrival in the Canary Islands places it close to mainland Europe. In this study, we explored how widespread the parasite is on Tenerife and what environmental conditions allow it to thrive. We examined rats, snails, and lizards from different parts of the island and found that the parasite is well established in all three groups. By combining these field data with environmental information, we built models to predict where the parasite is most likely to occur and compared Tenerife’s climate to Hawaii, where human infections are common. Our results show that suitable conditions exist across much of the Canary Islands, but areas where people live densely overlap only slightly with the parasite’s hotspots. This may explain why no human cases have been recorded so far, even though the parasite is abundant in wildlife.

## Introduction

*Angiostrongylus cantonensis* (commonly known as the rat lungworm) is an invasive parasitic nematode originally endemic to Southeast Asia. Since its first description in the 1930s in China [1], the species has exhibited a remarkable capacity for global dispersal. Over the following decades, its range has steadily expanded, and it is now present on all continents except Antarctica. Recent records suggest that the parasite is approaching or has already established new foci in or near Europe [2–5]. The rat lungworm is considered a highly successful invader, mainly due to the widespread distribution and invasive potential of its two primary host groups: rats (which serve as definitive hosts) and gastropods (intermediate hosts). Some representatives of both of these host groups are globally invasive and thrive in a wide range of environments far beyond their native ranges [6–10]. This close ecological association with other successful invaders makes *A. cantonensis* a striking example of parallel biological invasion, whereby the parasite expands its range in tandem with the global spread of its hosts.

The rat lungworm is not only an ecologically successful invader but also an important zoonotic pathogen. In humans, it is the primary cause of eosinophilic meningitis, which affects the central nervous system and is increasingly recognized as a new serious infectious disease. Clinical manifestations range from mild, flu-like symptoms to severe neurological impairment, and in rare cases, the infection can lead to death [11]. In addition to humans, *A. cantonensis* can infect a wide range of accidental or aberrant hosts, including other mammals and birds, in which it may cause neurological symptoms of varying severity. [12, 13].

Interestingly, despite the presence of the parasite in several regions of Europe, confirmed human cases are virtually absent or extremely rare. To date, only a single likely autochthonous case of neuroangiostrongyliasis has been reported in France [14]. This is in stark contrast to other affected regions, such as parts of the Pacific where human cases are relatively frequent [15]. One possible explanation lies in differences in cultural and dietary practices. In many endemic regions of Southeast Asia and the Pacific, raw or undercooked gastropods and paratenic hosts - such as freshwater shrimp, frogs, or monitor lizards - are sometimes consumed as part of traditional cuisine, increasing the risk of infection. In contrast, such practices are rare in Europe. Nevertheless, transmission can also occur through ingestion of contaminated vegetables, fruits, or drinking water, which may harbor infective larvae. This highlights the importance of food hygiene and public awareness even in regions where risky culinary practices are uncommon.

In addition to the newly recognized hotspots within Europe [3, 5, 16], the Canary Islands represent an important and relatively well-studied focus of the infection. The rat lungworm has been reported in Tenerife from its definitive hosts in 2010 [17] and has been studied quite extensively in its intermediate and paratenic hosts [4, 18–20]. Although the nematode has been relatively well studied in the region, significant knowledge gaps remain - particularly in predicting its probable distribution across the archipelago - which limits the implementation of targeted surveillance. Furthermore, current knowledge does not allow for the clear identification of key factors influencing the presence and prevalence of *A. cantonensis* throughout the islands. Interestingly, in contrast to other hotspots of rat lungworm infection with similarly high prevalence in intermediate and paratenic hosts [21–23] - which can serve as sources of human infection - not a single human case has been reported in the archipelago. While underdiagnosis cannot be entirely ruled out, the Canary Islands are a well-developed region with a relatively high standard of healthcare, making this explanation less likely.

To shed more light on the above mentioned matter, we present a comprehensive analysis of rat lungworm occurrence across three different host groups. In particular, we modelled spatial patterns in the distribution and prevalence of *A. cantonensis* in the Canary Islands and assessed relative importance of environmental factors driving these patterns. Our results highlight areas of high predicted presence that overlap with regions of dense human population. In addition, we provide a comparison of climatic conditions between the Canary Islands and Hawaii - another archipelago where *A. cantonensis* has been introduced and studied in recent years, but relatively numerous human cases have been reported [21].

Since Tenerife (and the Canary Islands in general) is characterized by diverse landscapes, a wide range of vegetation types, and substantial altitudinal variation (Fig 1), we hypothesize that the presence and prevalence of the nematode will vary considerably across the island. To date, relatively high prevalences have been reported in the northeastern tip of Tenerife [18, 19], while only a single positive sample in the southern part of the island was recently documented in a study focused on endemic lizards [4]. The apparent spatial disparity may reflect both environmental influences and uneven sampling effort, as host collection is generally easier in the humid north. Because existing data do not allow these factors to be disentangled, this study aims to fill this gap.

**Fig 1.**
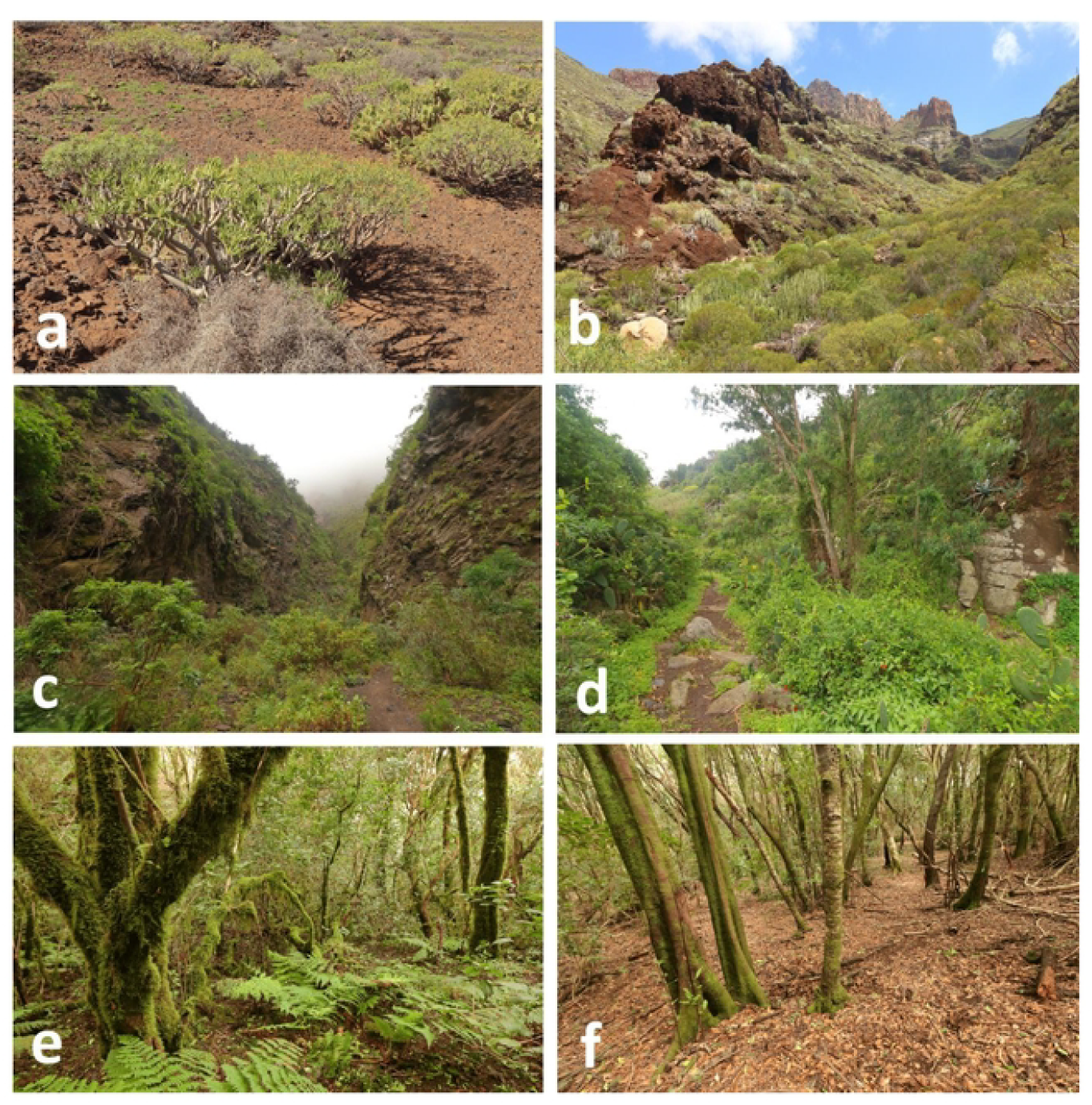
Habitats pictures of sampling sites. **a**,**b** Xeric vegetation types in Tenerife. (a) Bare volcanic rocks with scattered succulents. (b) Sparse xerophytic bushland. **c**,**d** Bush vegetation in Tenerife. (a) Continuous bush without tree cover. (b) Continuous bush interspersed with isolated trees. **e**,**f** Laurel forest in Anaga, Tenerife. Representative habitat photograph taken at a sampling locality. Photographs taken at sampling localities included in this study.

## Materials and methods

### Sample Collection

In total, we collected three host groups across 41 localities throughout the island, aiming to cover a range of altitudinal and vegetation zones with varying levels of anthropogenic influence (Fig 2). An area of maximum 400 m^2^ was considered one locality. All ethical requirements were met, and animal captures were authorized by the Canary Government in accordance with Law 42/2007, under expedient numbers 2022/14555 and 2022/11052. The capture methods for all species are described in detail below. Collection of lizard samples used for modelling is described in Anettová et al. (2024) [4]; preferably, only tail tissue was used, and the lizards were subsequently released.

**Fig 2.**
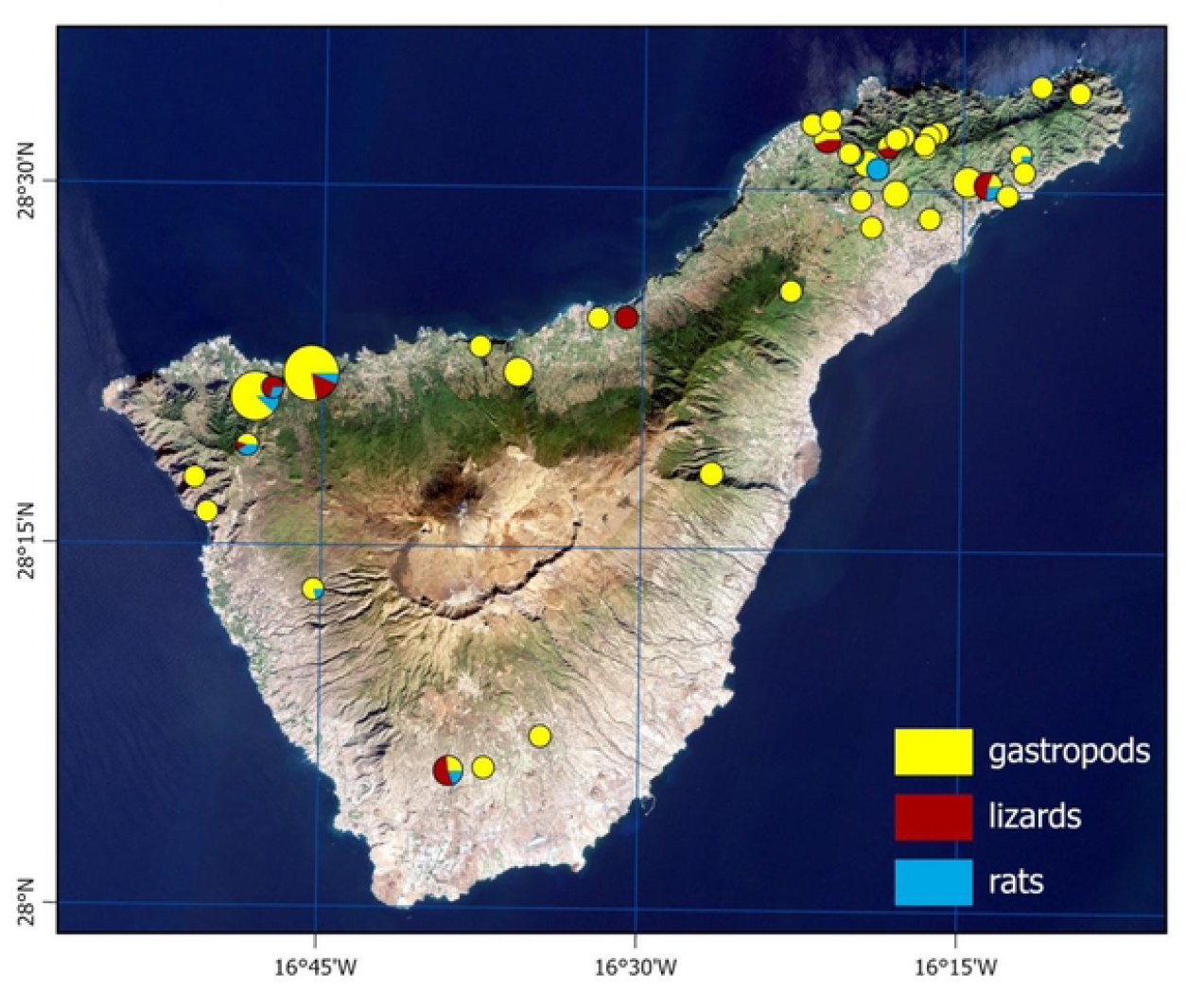
Sampling localities of gastropods, lizards, and rats across Tenerife. The size of the pie chart indicates the total number samples collected at each locality. Individual slices indicate the proportion of each group.

#### Rats collection

We collected definitive hosts, rats, at 11 localities. Animals were captured in live traps set at dawn, left overnight, and retrieved by dusk the following morning. Rats were euthanized using medetomidine (0.2 mg/kg) and ketamine (3.5 mg/kg), followed by injection of T61 intracardially (1 ml pro toto) by a person authorized to handle and euthanize experimental animals. Rats were subsequently dissected, and adult *A. cantonensis* nematodes were collected from the heart or pulmonary arteries. Brain samples were collected as well and all the samples were stored in 70% ethanol for the following molecular analysis. Individuals were considered positive if adult nematodes were found in their lungs or heart.

#### Gastropods collection

Gastropods were collected at 37 sites over the island. Each site, an area of approximately 20×20 m of homogeneous habitat, was surveyed for approximately 60 minutes. Gastropods, both native and non-native species, were collected by a handpicking method. Litter sieving was not used as it captures mostly minute species, which do not fit the scope of our study. Collected gastropods were euthanized by freezing and subsequently preserved in molecular-grade 99% ethanol. The identification of species was performed later in the laboratory. The nomenclature follows MolluscaBase eds. (2025).

### Molecular analyses

Out of all collected gastropods, 697 individuals were selected for molecular analysis. DNA was extracted using the DNeasy Blood & Tissue Kit (Qiagen, Germany), with an extended overnight pre-lysis step. For gastropods, approximately 25 µg of foot tissue was used, while a similar amount of cerebral tissue was processed in the case of rats. DNA of *A. cantonensis* adults was extracted by using approximately one third (middle part) of the nematode, following the same procedures as described above.

Quantitative PCR analysis was done using LightCycler 480, following protocol by Sears et al. [24]. DNA from a single third-stage larva of*A. cantonensis* extracted by the same method as the samples and diluted 100× was used as a positive control. Nuclease-free water was used as a negative control. The assay was run in duplicates. The Ct value (average value between duplicates) of positive samples was calculated by absolute quantification analysis of the 2nd Derivative Maximum. Only amplification curves with a Ct value under 35 were considered positive to avoid false positive results due to amplification and fluorescence artifacts, or cross-contamination.

In case of adult nematodes, DNA was extracted from approximately one third, middle part, of the adults’ body using innuPREP Forensic Kit (Innuscreen GmbH, Germany). Complete gene for subunit I of the cytochrome oxidase (COI) was amplified as described elsewhere [5]. PCR products were separated and visualised by an agarose gel electrophoresis and subsequently purified with ExoSAP-IT™ (Thermo Fisher Scientific, USA). Sanger sequencing was performed commercially in SEQme, s.r.o. (Czech Republic). Sequences were checked, trimmed and aligned in Geneious Prime 2025.2.2 (www.geneious.com) and their identity was checked against the GenBank database using BLASTn [25].

### Species Distribution Modelling

To model the potential distribution of *A. cantonensis* in the Canary Islands, we used presence data derived from the field surveys. Localities were considered positive if the parasite’s DNA was detected by molecular methods in any host species (gastropods, rats, or lizards), or if adult nematodes were found in the lungs or heart of rats.

Species distribution modelling was performed using the Maximum Entropy algorithm implemented in MaxEnt software (version 3.4.4). Environmental predictor variables included bioclimatic data, land cover, vegetation indices, and topographic characteristics (Supplementary Table 1). All raster layers were resampled to a common resolution of approximately 100 *×* 100 m (0.0012555749°) and kept in WGS 1984 coordinate system to match the coordinate format of the occurrence data. To ensure consistent model calibration focused on Tenerife, a bias file was used to constrain background point selection to this island.

The bioclimatic variables were obtained from the CHELSA database (for period 1981–2010 [26]) and included: annual mean temperature (bio1), temperature seasonality (bio4), annual precipitation (bio12), and precipitation seasonality (bio15). These were originally at 1 km resolution and resampled accordingly. Landscape structure was represented by a 2018 Tree Cover Density (TCD) layer from the European Union’s Copernicus Land Monitoring Service, and six CORINE Land Cover (CLC) categories transformed into binary layers. These categories included urban areas (class 1), agricultural land (classes 21, 22 and 24), forests (class 31), scrub vegetation (class 32), pastures and grasslands (classes 231 and 321) and non-vegetated areas (class 33).

Local site moisture conditions were represented by a Topographic Wetness Index (TWI), calculated in SAGA GIS (version 8.3.0) based on the Copernicus Global DEM at 90 m resolution [27]. Vegetation dynamics were incorporated via mean NDVI and NDVI standard deviation for the year 2022 [28]. Terrain Ruggedness Index (TRI) was ∼ also included as a topographic predictor. All raster layers were resampled to a common resolution of approximately 100 × 100 m (∼0.00126°) using a B-Spline interpolation in SAGA GIS.

Models were trained with default MaxEnt settings unless stated otherwise. Variables contributing less than 0.5% to the model were excluded to improve model parsimony. The final model was projected to all Canary Islands to identify suitable habitats beyond Tenerife.

Prevalence of *A. cantonensis* was calculated for 37 localities in Tenerife as the proportion of positive samples (min = 0, mean = 0.280, max = 0.7). Localities with fewer than six samples or samples marked as “failed” or “dubious” were excluded. To model the prevalence, we used the Boosted Regression Tree (BRT) algorithm and the same set of environmental predictors as in the MaxEnt modelling.

To assess the overlap between areas with high density of human population and predicted suitability for *A. cantonensis*, we overlaid the MaxEnt and the prevalence models output with land cover data on urban areas from CORINE Land Cover (class 1).

The BRT model was run in R using the *gbm*.*step* function from the *dismo* package. We used the following parameters: family = “gaussian”, tree.complexity = 3, learning.rate = 0.001, bag.fraction = 0.75, and step.size = 10. The optimal number of trees was selected via 5-fold cross-validation. Localities were weighted by the number of host samples collected per site (min = 6, mean = 23.84, max = 106). An initial model was run with all predictors, and those with zero contribution were subsequently removed before refitting the final model using the same settings.

To assess climatic similarity between the Canary Islands and Hawaii (area with known human cases of *A. cantonensis* infection), we performed a Multivariate Environmental Similarity Surface (MESS) analysis in R using the *dismo* and *raster* packages [29]. The analysis used the following bioclimatic variables: *bio1*, bio4, bio12, and bio15, as described above. MESS was trained on climatic data from Hawaii, then projected to the Canary Islands to identify areas of environmental similarity and novelty relative to the training region.

## Results

### *Angiostrongylus cantonensis* is widely distributed across Tenerife

Out of 41 surveyed localities, 35 were confirmed positive based on at least one host species (gastropods, rats, or lizards). In total, 78 rats (both *R. rattus* and *R. norvegicus*) were collected from 11 sites, of which 13 tested positive (16.7%). The highest prevalence in rats was observed in Tegueste (village in the humid northeast of the island), where half of the examined individuals carried the parasite. All adult nematodes collected from rats were confirmed as *A. cantonensis*, and their COI sequences were identical across all specimens (GenBank accession number: PX496382). Among reptiles, 129 lizards (*Gallotia galloti*) were examined, with 35 positives (27.1%) [4]. Anaga (national park in the humid northeast) was the site with the highest infection rate of lizards, with a prevalence of 63.6%. Gastropods represented the largest sample set: 697 individuals from 37 localities were tested, of which 185 (26.7%) were positive and four were inconclusive. Again, the Anaga region showed the highest prevalence in gastropods (43.3%). After excluding sites with fewer than six samples or with failed/dubious results, prevalence estimates were calculated for 37 sites and ranged from 0 to 0.70 (mean = 0.28).

### Models predicted the highest environmental suitability in the humid northeast of Tenerife

MaxEnt predictions showed the highest habitat suitability in the northeastern tip of the island, with scattered patches in mid-altitude zones. There were eight variables retained in the final model: precipitation seasonality, annual mean temperature, topographic wetness, terrain ruggedness, tree cover density, vegetation indices and temperature seasonality. The most influential predictors were precipitation seasonality (percent contribution 52.1%), annual mean temperature (18.1%), and terrain ruggedness (9.5%) (Table 1). Response curves indicated that the parasite favored intermediate temperatures and lower rainfall variability (Supplement Fig 1).

**Table 1.** Percent contribution and permutation importance of environmental variables retained in the MaxEnt model.

| Variable | Percent contribution | Permutation importance |
| --- | --- | --- |
| bio15 (Precipitation seasonality) | 52.1 | 9.6 |
| bio1 (Annual mean temperature) | 18.1 | 49.8 |
| SAGA TWI (Topographic Wetness Index) | 10.4 | 9.4 |
| TRI (Terrain Ruggedness Index) | 9.5 | 20.3 |
| TCD 2018 (Tree Cover Density) | 8.4 | 0.9 |
| NDVI mean (2022) | 0.8 | 1.6 |
| NDVI SD (2022) | 0.5 | 0.4 |
| bio4 (Temperature seasonality) | 0.4 | 7.9 |

Intersecting the MaxEnt prediction with urbanised areas showed that the overlap between highly suitable habitats and densely urbanised zones (i.e., areas of high population density) was limited, mainly restricted to small patches in the northeastern outcrop (Fig 3).

**Fig 3.**
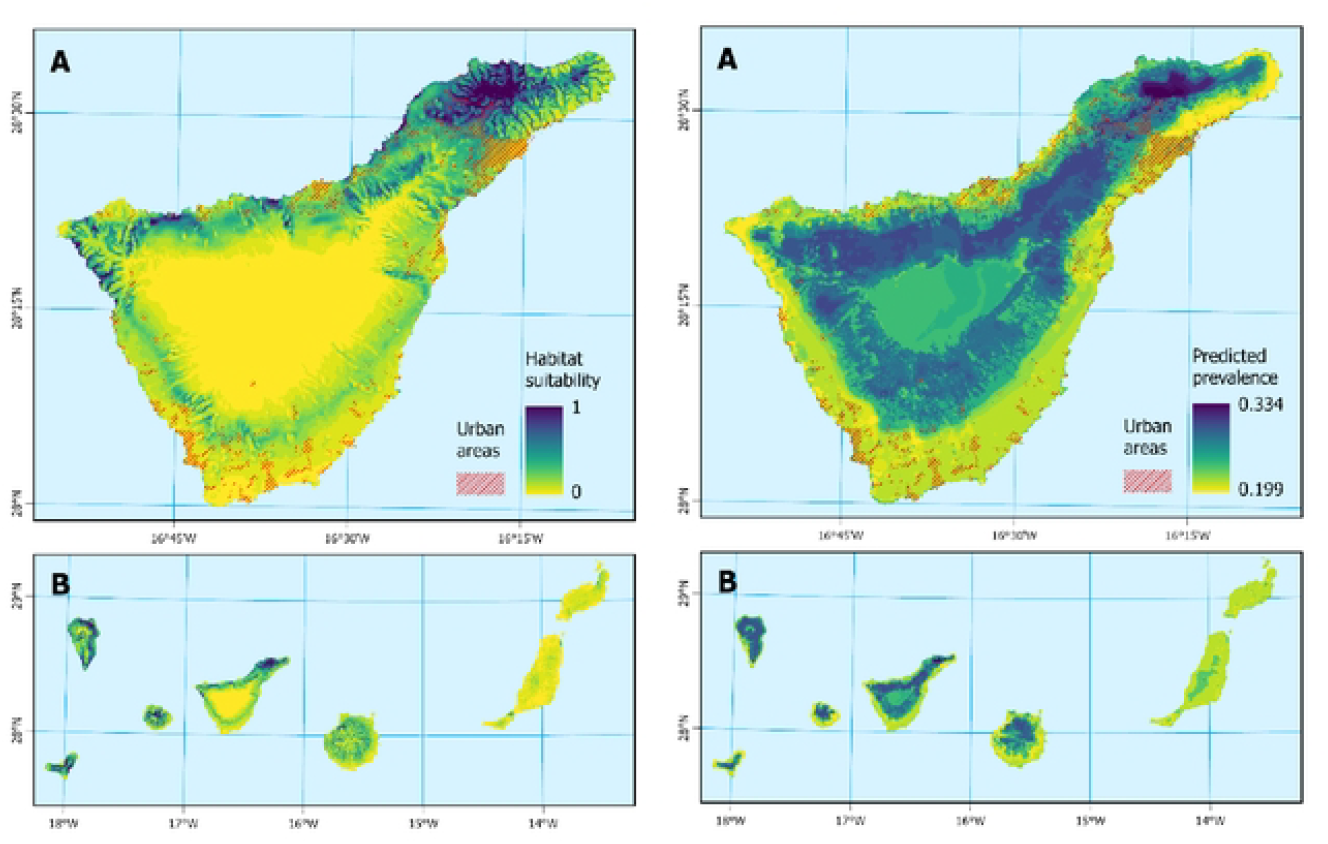
MaxEnt model prediction of suitable habitat for *A. cantonensis* (left), with the overlap of urban areas (CORINE land cover, class 1). (a) Detailed prediction for Tenerife. (b) Predicted prevalence across the Canary Islands; and **b** Boosted Regression Tree (BRT) model prediction of *A. cantonensis* prevalence (right). (a) Detailed prediction for Tenerife. (b) Predicted prevalence across the Canary Islands.

### Landscape structure and climate strongly influenced parasite prevalence

The BRT model (pseudo-*R*^2^ = 26%, MSE = 0.0412) confirmed that tree cover density (30.9%), precipitation seasonality (22.2%), and temperature seasonality (17.0%) were the strongest predictors of prevalence variation among sites. Vegetation indices also contributed notably (NDVI mean = 12.7%, NDVI variability = 6.9%). In contrast, annual precipitation and topographic wetness had minimal influence (*<*1%). The BRT model predicted prevalence values ranging from 20% to 32% across Tenerife. Predicted prevalence maps highlighted foci overlapping with the northeast, but also indicated suitable mid-elevation habitats outside the current known distribution (Fig 3). Detailed contributions of the variables are shown in (Table 2).

**Table 2.** Relative influence of environmental variables in the BRT model.

| Variable | Relative influence (%) |
| --- | --- |
| Tree cover density | 30.95 |
| Precipitation seasonality | 22.19 |
| Temperature seasonality | 17.03 |
| NDVI mean | 12.74 |
| NDVI standard deviation | 6.88 |
| Mean annual temperature | 5.83 |
| Terrain ruggedness index (TRI) | 3.21 |
| Annual precipitation | 0.79 |
| Topographic wetness index | 0.36 |

### Dry leeward zones shared climatic conditions with Hawaii, whereas the humid northeast was novel

MESS projections using bioclimatic variables (bio1, bio4, bio12, bio15) indicated relatively high environmental similarity in the drier leeward areas - southern Tenerife, southern Gran Canaria, and El Hierro - while the humid northeast of Tenerife showed low similarity (i.e., novel conditions relative to Hawaii, Fig 4). This suggests that transmission dynamics inferred from Hawaiian systems may translate primarily to drier Canary Island habitats, whereas the humid northeastern sector may follow its own unique pathways.

**Fig 4.**
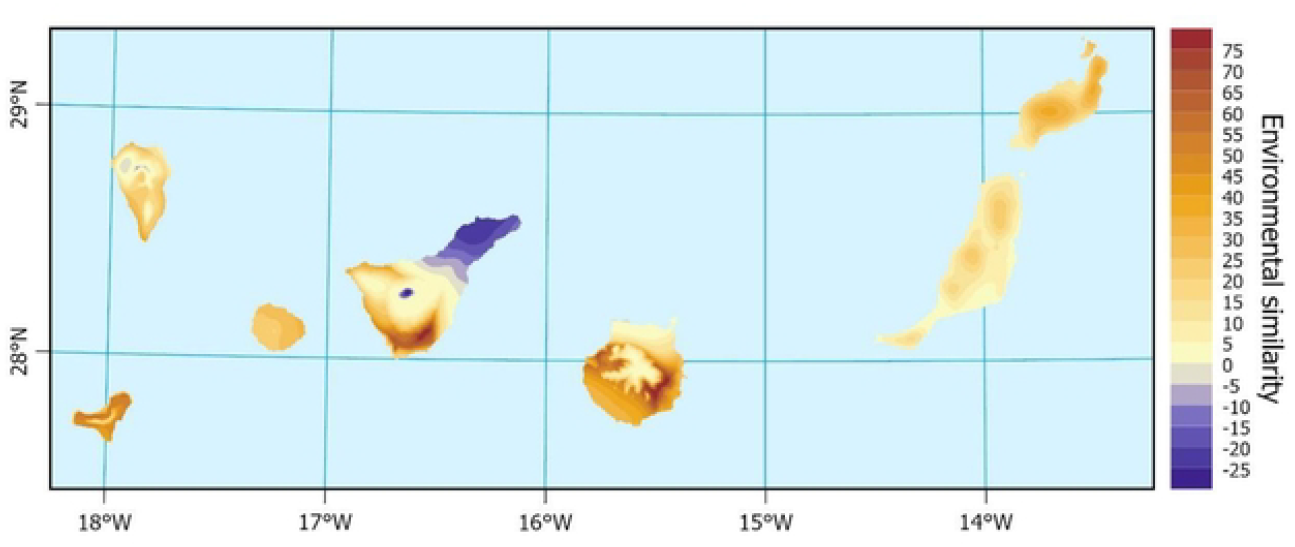
Multivariate Environmental Similarity Surface (MESS) analysis. Model trained on climatic data from Hawaii and projected onto the Canary Islands using selected bioclimatic variables.

## Discussion

Our survey confirms that *A. cantonensis* is now firmly established across Tenerife. The parasite was detected in all parts of the island, with 35 of 41 surveyed localities testing positive, including arid southern sites where it had been previously reported in endemic lizards [4]. The invasion process is typically described as a sequence of five stages: transport, introduction, establishment, spread, and negative impacts [30].

Establishment can be recognized by the successful exploitation of local hosts and an increasing number of infected host species over time [31, 32]. Since the parasite was first recorded on Tenerife in 2010 [17], it has circulated among both endemic and invasive species, involving intermediate as well as paratenic hosts [18, 20]. We therefore consider *A. cantonensis* to be well established on Tenerife. The situation on the remaining Canary Islands is less clear, although its presence has been documented [33]. The broad host range raises concerns not only for zoonotic risk but also for potential impacts on wildlife, including birds and other accidental hosts that may suffer from neurological disease.

Despite this well-documented presence in animal hosts, no human neuroangiostrongyliasis cases have been reported from Tenerife, mirroring a broader pattern across Europe where the parasite is present but human cases are absent or extremely rare [3, 34]. One plausible explanation lies in local cultural and culinary practices. In contrast to Southeast Asia and parts of the Pacific, where raw or undercooked gastropods, frogs, or lizards are often consumed, such practices are not common in Europe. Sporadic cases cannot be entirely ruled out, but the absence of outbreaks suggests that dietary habits and food preparation standards strongly limit the risk of human infection.

In Tenerife specifically, our models highlight another potential factor: the overlap between areas of highest predicted environmental suitability for *A. cantonensis* and densely populated urban zones was limited, restricted to small patches in the humid northeast. Most of the suitable habitat is instead associated with forested or semi-natural areas, where human–gastropod contact is expected to be less frequent. The parasite’s dependence on precipitation seasonality and tree cover density, as identified by MaxEnt and BRT analyses, reinforces this pattern, as these conditions are most pronounced in natural and forested landscapes rather than in croplands or densely urbanised areas.

It is important to note that the BRT model, explaining only 26% of variation in prevalence values, produced predictions in a narrow range of 20–32% rather than across the full 0–70% prevalence spectrum. Although the model was not able to predict exceptionally high prevalences, the spatial pattern was well captured, confirming the results of Maxent modelling.

The MESS analysis further underscores that climatic similarity to regions with known human cases varies across the Canary Islands. Drier leeward zones of Tenerife, southern Gran Canaria, and El Hierro share relatively high climatic similarity with Hawaii, whereas the humid northeastern tip of Tenerife, where prevalence is highest, represents novel conditions outside the Hawaiian reference climate conditions. This apparent paradox - high prevalence but low climatic analogy to Hawaii - illustrates that bioclimatic factors alone cannot explain or predict human infection risk, probably not even the presence of the parasite by itself. Rather, it highlights the need to distinguish between two levels of the problem: first, the establishment and circulation of the parasite in its natural hosts, which is shaped by climate, invasion history, and rat ecology; and second, the emergence of human cases, which depends far more on ecological, cultural, and socioeconomic drivers of exposure.

Comparisons with other regions support this view. In Hawaii, where the parasite was introduced and is now widespread, human cases are frequent [21] despite a largely Western-style diet, likely due to anthropological and ecological factors such as the use of rainwater catchment systems and inadvertent consumption of contaminated produce. In Australia, meanwhile, cases of neuroangiostrongyliasis are more commonly reported in dogs [35], again reflecting local exposure pathways rather than climate alone. These examples suggest that the parasite’s broad ecological tolerance and ability to persist in rats and snails effectively decouple its distribution from strict climatic boundaries. Its capacity to overwinter in rodents due to long patent periods, and to persist in snails as shown experimentally [36], further widens the climatic niche beyond what environmental predictors can capture.

Taken together, our results highlight that species distribution models are highly useful for identifying environmental hotspots and guiding surveillance, but they alone cannot predict where human infections will occur. The absence of human cases in Tenerife and Europe more broadly, despite established host cycles, points towards exposure-related factors as the critical determinant. Future research should therefore aim to integrate ecological modelling with anthropological and epidemiological perspectives, focusing on dietary practices, food safety, water systems, and other human behaviors that modulate the parasite’s zoonotic potential.

## Conclusion

Our study shows that *A. cantonensis* is widely established across Tenerife, infecting definitive (rats), intermediate (gastropods), and paratenic (lizards) hosts, with prevalence exceeding 60% in some localities. Species distribution modelling identified the humid northeastern part of the island as the main hotspot, with tree cover, temperature, and rainfall stability as the strongest environmental drivers of occurrence and prevalence. Although suitable habitats extend into urbanised zones, their overlap with dense human settlements was limited, which may explain the current absence of reported human cases. These findings highlight the successful establishment of the parasite in Tenerife and underscore the need for continued surveillance at the wildlife-human interface.

## Authors’ contribution

Investigation: L.A., R.C., A.Š., V.B., J.K., E.I.R., B.Č.; Writing – review & editing: J.D., R.C., M.H., D.M.; Methodology: L.A., A.Š., V.B., J.D., D.M., L.A.; Visualization: L.A., J.D., R.C.; Conceptualization: D.M., L.A., M.H.; Data curation: D.M., J.D., M.H.; Supervision: J.D., M.H., D.M.; Validation: J.D., M.H., D.M.; Writing – original draft: L.A.; Funding acquisition: D.M.; Project administration: P.F.; Resources: P.F.

## Ethical standard

The authors declare that all ethical standards and requirements were fulfilled. Animal trapping and use was approved by the Canary Government in accordance with Law 42/2007 with the expedient numbers 2021/29507 and 2022/11052.

---

**Supplemnt Figure 1.**
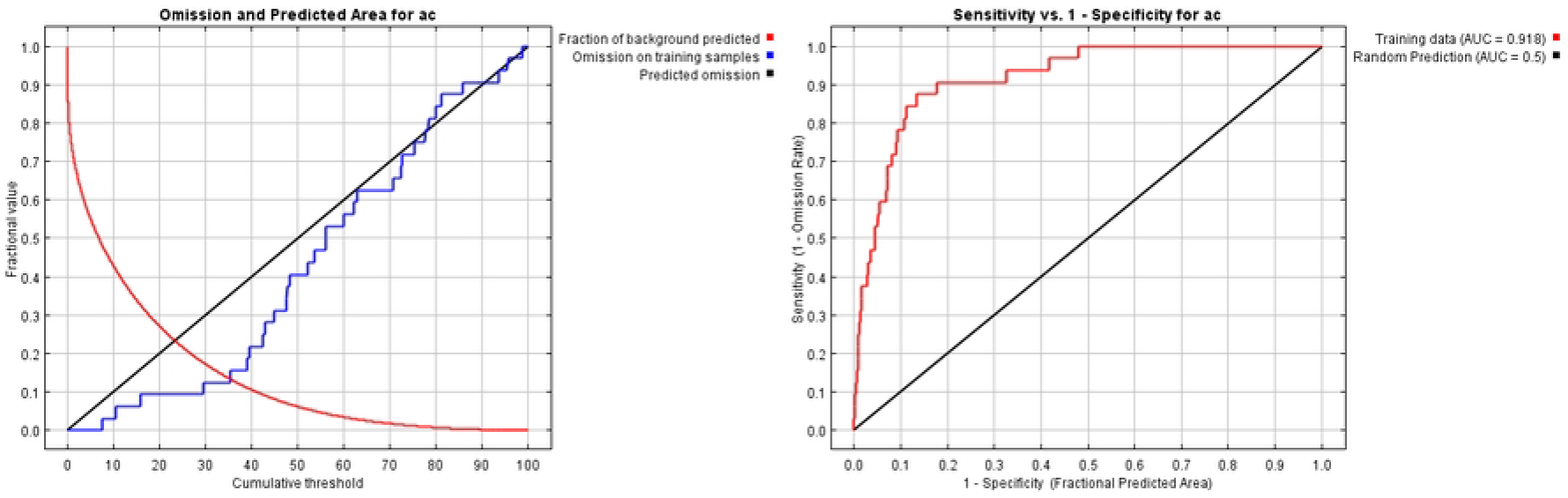
Box initially at rest on sled sliding across ice.

**Supplemnt Figure 2.**
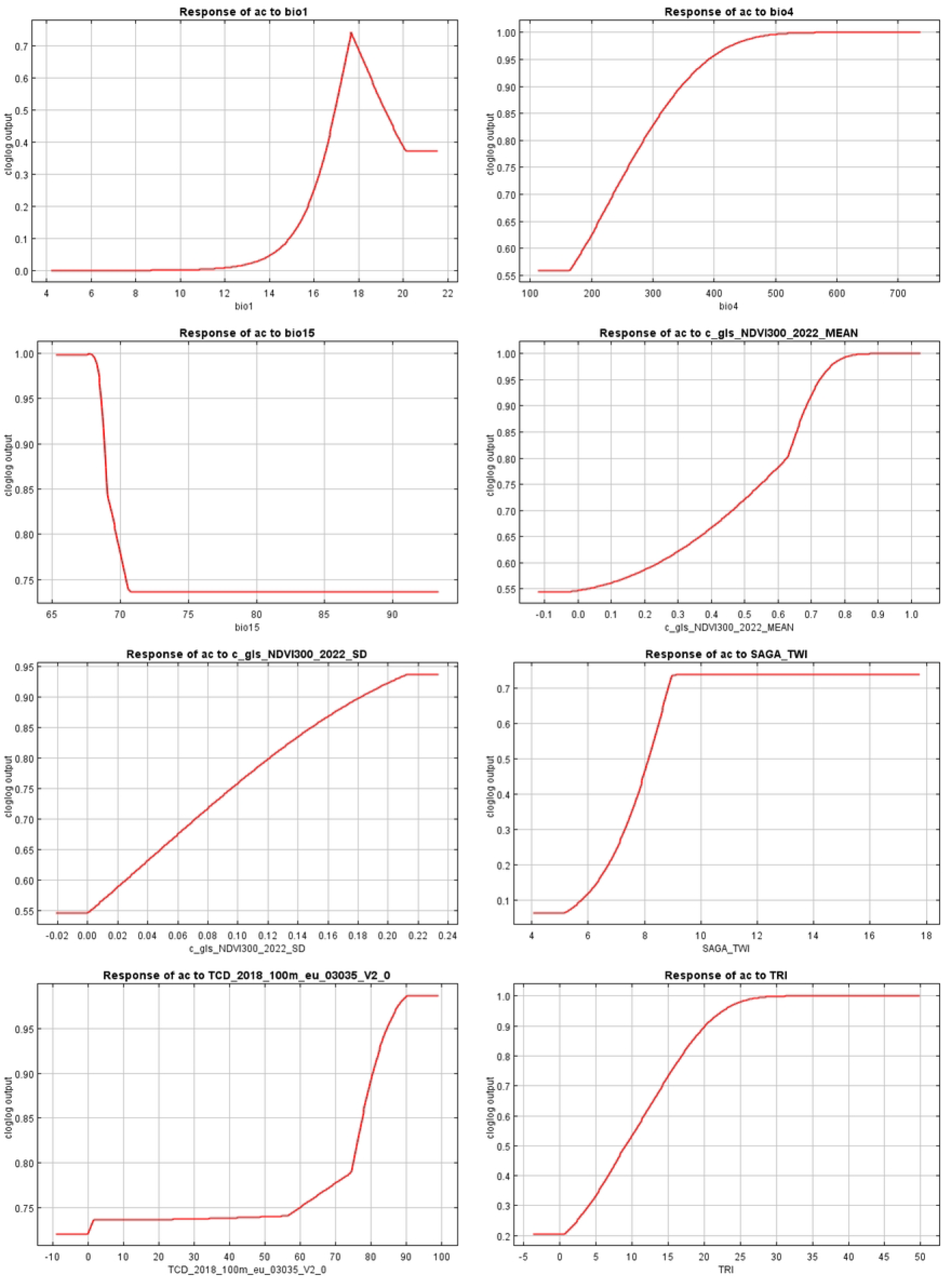
Box initially at rest on sled sliding across ice.

